# Acute radiofrequency electromagnetic radiation exposures cause neuronal DNA damage and impair neurogenesis in the young adolescent rat brain

**DOI:** 10.1101/2020.11.07.370627

**Authors:** Kumari Vandana Singh, Chandra Prakash, Jay Prakash Nirala, Ranjan Kumar Nanda, Paulraj Rajamani

## Abstract

Mobile phone is now a commonly used communication device in all age groups. Young adolescents use it for longer duration and effect of its radiofrequency electromagnetic radiation (RF-EMR) on their brain structure and function need detailed investigation. In the present study, we investigated the effect of RF-EMR emitted from mobile phones, on young adolescent rat brain. Wistar rats (5 weeks, male) were exposed to RF-EMR signal (2,115 MHz) from a mobile phone at a whole body averaged specific absorption rate (SAR) of 1.15 W/kg continuously for 8 h. Higher level of lipid peroxidation, carbon centered lipid radicals and DNA damage were observed in the brain of rat exposed to RF-EMR. Number of neural progenitor cells (NPCs) in dentate gyrus (DG) were found to be relatively low in RF-EMR exposed rats that may be due to reduced neurogenesis. Acute exposure to RF-EMR induced neuronal degeneration in DG region with insignificant variation in CA3, CA1 and cerebral cortex sub regions of hippocampus. Findings of this study, indicate that acute exposure of high frequency RF-EMR at relatively higher SAR may adversely impact the neurogenesis and function of adolescent rat brain. Generation of carbon centered lipid radicals, and nuclear DNA damage might be playing critical role in reduced neurogenesis and higher neuronal degeneration in the cortex and hippocampus of brain. Detailed understanding of RF-EMR induced alteration in brain function will be useful to develop appropriate interventions for reducing the impact caused by RF-EMR damage.

## 1. Introduction

Approximately 6.9 billion people are using mobile phones worldwide and substantial fraction of it (3.8 billion) are using smart phones (www.statistica.com). Smart phones with third generation technology (3G) operates at a frequency range of 1.8-2.5 GHz without periodic pulsed modulation and phones with fourth generation (4G) technology, operates at a higher frequency range of 2-8 GHz with Long-term Evolution (LTE) modulation (Kesari et al. 2014; Miller et al., 2019). Growing body of evidence from epidemiological studies suggest that use of smartphones among children and adolescent is associated with behavioral and emotional disorders (De-Sola Gutierrez et al., 2016; Foerster et al., 2018; Sudan et al., 2016). Generation of oxidative stress, epigenetic changes and DNA damage are seemed to be underlying mechanisms for adverse effects of radiofrequency electromagnetic radiation (RF-EMR) exposure on developing brain (www.bioinitiative.org; Sage and Burgio, 2018). However, the molecular mechanism causing these effects on young brain needs additional studies.

Owing to their thin skull, lower myelin content and higher water content, young brain absorbs 2-3 times higher dose of RF-EMR energy emitted from mobile phones placed against head compared to adults (Fernandez et al., 2018; Redmayne and Johansson, 2014). In addition, many developmental changes and ongoing maturational process in young adolescent brain makes them susceptible to RF-EMR (Belpomme et al., 2018). Cerebral cortex and hippocampus absorb maximum RF-EMR energy at higher frequencies (Cardis et al., 2008; Fernandez et al., 2018). Earlier studies showed RF-EMR induced adverse effects on developing rodents largely depends on the frequency and specific absorption rate (SAR) of radiation exposure (Chen et al., 2014). Further, dentate gyrus (DG) region of the hippocampus is the site of adult neurogenesis and play important role in memory acquisition and highly sensitive to early life stressors (Huang, 2014; Youssef, et al., 2019). So, the implications of higher RF-EMR energy absorption in these brain regions of young brain need focused studies.

SAR is an important determinant for thermal effect of RF-EMR and other factors like frequency, duration of exposure and modulation as well as physiological parameters and developmental stage of animal modulate their biological effects (Belyaev, 2010; IARC, 2013). Like, certain oscillatory frequencies of non-thermal RF-EMR may interfere with endogenous electric field of nervous system (Weinwright, 2000), which is particularly detrimental for proliferation and differentiation of neurons in young brain (Levin, 2003). Incidences of glioma has been reported in the actively dividing glial cells of young brain when exposed to RF-EMR without causing any significant tissue heating (Wyde et al., 2016; Falcioni et al., 2018).

Several reports demonstrated detrimental effect of non-thermal RF-REMR on hippocampal pyramidal, or cortical neurons of young rats leading to learning and memory impairment (Bas et al., 2009; Deshmukh et al., 2015; Kumari et al., 2017). In vitro and in vivo studies have shown that long term exposure of mobile phone radiations causes oxidative stress in the brain (Yakymenko et al., 2016). RF-EMR induced oxidative damage to cellular DNA has shown mixed results (www.bioinitiative.org; Belyaev, 2015). Effect of RF-EMR exposure on adult hippocampal neurogenesis need additional investigation. Previously, few studies have investigated the effect RF-EMR exposure on neurogenesis with mixed findings (Kim et al., 2015; Odaci et al., 2008; Xu et al., 2017). Electromagnetic field reported to epigenetically interferes with proliferation of neural progenitor cells (NPCs) in adult tissue through reactive oxygen species (ROS) production (Falone et al, 2007; Park et al., 2014).

Majority of the studies have focused on undertaking long term consequences of RF-EMR exposure. These studies have limitation to miss damage partly corrected or adopted by the organism and may miss to characterize the early loss of function. Therefore, in this study we employed a comprehensive approach to estimate oxidative stress, DNA damage and neuronal degeneration in cortex and hippocampus regions of brain as well as proliferation of hippocampal NPCs in male Wistar rats of age comparable to young adolescents which are subjected to RF-EMR in a exposure set up mimicking short-term acute exposure (8 h continuously) from a smartphone (2115 MHz) operating at higher SAR value (whole body average SAR value of 1.1 W/kg).

## 2. Methodology

### 2.1 Animals and experimental plan

Procedures approved by the institutional animal ethical committee (IAEC) of Jawaharlal Nehru University, New Delhi, India (Application code # 23/2016) were followed to carry out the animal experiments. Wistar rats (Male, 5weeks, n=26) were acquired from Central Laboratory Animal Resources, Jawaharlal Nehru University, New Delhi and housed under 12 hours light dark cycle at 25 °C room temperature with food and water provided *ad libitum.* After one week of habituation, animals were randomly divided in to sham and exposed groups and housed in the group of two animals per cage. A schematic presentation of the experimental plan adopted in this study is shown in **Fig. 1**.

**Figure 1.**
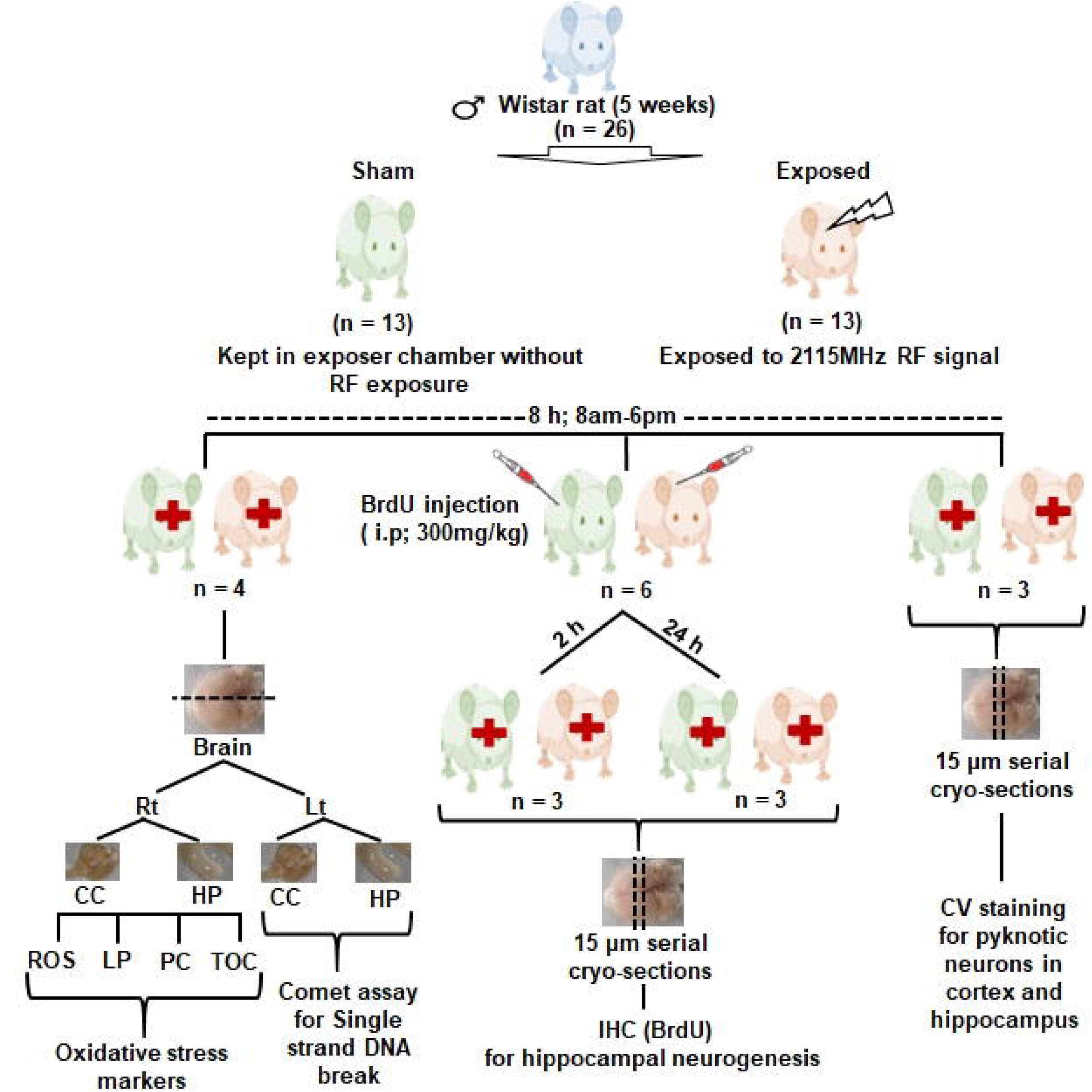
Workplan adopted to study the effect of radiofrequency electromagnetic radiation (RF-EMR) exposure for 8 h on developing brain of Wister rats. Total twenty-six rats were divided into two groups, 1) Sham and 2) Exposed. Out of thirteen rats from each group four were used to study oxidative stress parameters (one hemisphere) and comet assay (another hemisphere), and three rats for cresyl violet (CV) staining. Six rats from each group were injected with BrdU intraperitoneally (300 mg/kg) at the end of exposure time and sacrificed 2 h and 24 h post-injection (n=3) to study hippocampal neurogenesis. BrdU: 5’-bromo-2’-deox-yuridine, CC: cerebral cortex, HP: hippocampus, LP: lipid peroxidation, PC: protein carbonylation, ROS: reactive oxygen species, TOC: total antioxidant capacity.

### 2.2 Radiofrequency exposure

Study animals of exposed group were subjected to RF-EMR once for 8 h continuously between 6 a.m. to 2 p.m. Exposure set up consists of vector signal generator (VSG25A, Sigma Hound, USA) emitting 2115 MHz RF-EMR with digital modulation (16 QAM) through a horn antenna (10 cm diameter). The horn antenna was placed inside a specially designed anechoic chamber (1 m × 1 m ×1.5 m). During exposure experiment, animals were placed in Plexiglass rockets (5-7.5 cm × 20 cm × 6-8 cm) arranged in a circle inside the chamber with their truncated cones just below the center of horn antennae to allow equal exposure of all animals simultaneously (**Supplementary Fig. 1**). Perpendicular distance of the plexiglass rockets from antenna was kept at 6 cm to meet the nearfield exposure condition following the equation r ≤ 2D^2^/λ, where r is the distance between source of radiation and irradiated subject, D is the diameter of antenna, and λ is the wavelength of emitted RF-EMR. Each rocket is spacious enough to allow free body movement of animal and has holes (1 cm diameter) on upper and four sides to facilitate air flow. Receiving frequency and power at the level of rat body surface were measured using a monopole probe attach to spectrum analyzer (N9912A, Field fox, Agilent technology, USA). Power density was measured by the power meter (NBM-520, Narda, Germany) both outside and inside the cage during the exposure to adjust the fluctuation if any. Attenuation due to plexiglass sheet (2 mm) comes out to be 0.11 mW/cm^2^. So, power density at the level of cage (outer side) was kept at 1.11 mW/cm^2^ to maintain an effective power density of 1 mW/cm^2^ inside the cage. The whole-body SAR was estimated according to the equation explained in the earlier report (Durney et al., 1979). Following equation was used, which is an expression of average SAR for incident power density of 1 mW/cm^2^ and electric field (E) field polarization for a spheroid with length and width of animal taken as *semimajor* axis, a, and *semiminor* axis, b, in meters.

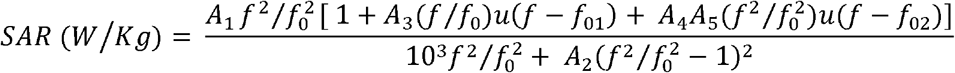

Where f_0_ is the resonance frequency, a function of a and b, f_01_ and f_02_ are empirically derived quantities depending on values of f_0_, a and b. u (f – f_01_) and u (f - f_02_) are the unit step functions. A1, A2, A3, and A4 are functions of a and b, and A5 is a function of ε, complex permittivity of muscle.

A group of study animals were kept in chamber under similar condition for same duration as exposed one, on the next day without RF-EMR exposure to include as Sham control. Out of total 26 animals, 13 were exposed to RF-EMR in the group of 4, 6 and 3 animals on alternate days (4 on 1^st^ day, 6 on 3^rd^ day, 3 on 5^th^ day) and their respective sham controls were kept in chamber on every alternate day (4 on 2^nd^ day, 6 on 4^th^ day, 3 on 6^th^ day). To minimize experimental variation, single exposure setup was used for all these experimental analyses. Exposure parameters were tabulated in **Supplementary Table 1**.

### 2.3 Preparation of tissue homogenate

Animals from both standing group (n=4) were sacrificed immediately after 8 hours of RF-EMR exposure using sodium pentobarbital (50 mg/kg, i.p.) followed by cervical dislocation. Whole brain was isolated followed by rapid dissection of hippocampus and cerebral cortex. Referring one hemisphere was used to study oxidative stress parameters and the other was used for comet assay. Tissue samples were flash frozen in liquid nitrogen and then stored at −80 °C until the experimental analysis. For oxidative stress analysis tissue was homogenized in ice cold phosphate buffered saline (PBS, 0.1 M pH 7.5) supplemented with Diethylenetriaminepentaacetic acid (DTPA; 2 mM) as transition metal chelator and protease inhibitor cocktail (P8340, Merck, USA) using mortal pestle at 4 °C. Homogenate was centrifuged at 1000 × g for 5 min at 4 °C to remove the debris and total protein concentration was estimated using Bradford reagent (5000006, Bio-Rad, USA).

For comet assay harvested tissues were minced using scalpel in 9:1 (v/v) mixture of Hanks’ Balanced Salt Solution (HBSS) media (14025092, Thermo Fisher scientific, USA) and DMSO. Next, the mixture was gently triturated with 5 ml pipette and allowed to settle for 1 min to remove debris. Single cell suspension thus obtained was used for the comet analysis. The entire procedure to obtain single cells was carried out within 5 min and at 4 °C to minimize DNA damage during experimental procedure.

### 2.4 Electron paramagnetic resonance (EPR) spectroscopy for detection of free radicals

Electron paramagnetic resonance (EPR) spectroscopy was performed to estimate concentration of carbon and oxygen centered free radicals using 5,5-Dimethyl-1-pyrroline N-oxide (DMPO) (Sigma-Aldrich, USA) a cyclic nitrone as a spin trap. Samples were prepared following the earlier published method (Rokyta et al. 2004). Briefly, tissue homogenate (54 μl) was mixed with DMPO (6 μl, 1 M) under nitrogen flow. EPR spectroscopy was performed using Bruker EMXmicroX-band (9.1-9.9 GHz) EPR spectrometer equipped with a Bruker ER 4102ST resonator/cavity maintained at 150 K using Bruker ER 4111 VT variable temperature unit. Sample mixture (30 μl) was immediately transferred to deep frozen X-band capillary tube (Supracil, Q-1.0X1.2 Wilmad-LabGlas, USA) and both ends were sealed. Capillary tube with sample was then immediately placed inside Bruker ER 169DIS quartz dewar in the EPR resonator in such a way that whole sample was vertically centered in the cavity. The spectra were recorded with following settings: central field; 3508.8 G, sweep width; 118.8 G, resolution; 1024 points, microwave frequency; 9.8 GHz, microwave power; 13 mW, modulation frequency; 100 kHz, modulation amplitude; 1.0 G, receiver gain; 2 × 10^3^, time constant; 163.8 ms, sweep time; 40.9 s, conversion time; 40 ms. Spectra of each sample was acquired and baseline corrected using WIN-EPR acquisition software. Reproducibility of the EPR signal was assessed in four study samples from each group. Signal intensities of radical species were determined as peak to peak height (arbitrary units) in first derivative spectrum which is a measure of radical concentration. Data was normalized with protein concentrations and expressed as arbitrary units (a.u.)/mg protein.

### 2.5 Lipid peroxidation assay

Malondialdehyde (MDA) concentration was determined following the method earlier reported (Uchiyama and Mihara, 1978). Briefly, tissue homogenate (125 μl) was mixed with Tris KCl Buffer (875 μl, pH 7.4) along with trichloroacetic acid (TCA; 30 %, 125 μl) and thiobarbituric acid (TBA, 52 mM, 125 μl). This solution was vortexed and incubated in water bath at 90 °C for 60 min and cooled in tap water. TBA reactive substances (TBARS) were extracted by centrifugation at 3,000 × g for 10 min. TBARS amount was determined by taking absorbance at 532 nm and was calculated as nanomole MDA equivalents per milligram of protein using the molar extinction coefficient of MDA (1.56 × 10^-5^ nM^-1^cm^-1^).

### 2.6 Protein oxidation assay

Protein oxidation in tissue homogenate was monitored following the earlier published method (Dubey et al., 1996). Briefly, the tissue homogenate (10 μl) was mixed with 2,4-Dinitrophenylhydrazine (DNPH, 40 μl, 10 nM) in HCl (2 M). Mixture was incubated for 1 h at room temperature in dark with continuous stirring and equal volume of TCA (20 %) was added with vortexing. Next, samples were kept on ice for 5 min and centrifuged at 10,000 × g for 10 min at 4 °C. Supernatant was discarded and pellet was washed once with TCA (10 %), and thrice in ethanol/ethyl acetate (1:1, v/v). Final precipitates were dissolved in guanidine hydrochloride (6 M) solution. Simultaneously, a blank was prepared by adding HCl (2 M, 40 μl) alone instead of DNPH. The difference (□) in absorbance of each sample was determined with reference to the blank reading at 370 nm. The amount of protein carbonyl was calculated using molar extinction coefficient of DNPH (2.2 × 10^-5^ nM^-1^cm^-1^) and results were expressed as nanomole carbonyls per milligram of protein.

### 2.7 Total antioxidant capacity assay

Total antioxidant capacity of tissue homogenate was estimated by measuring the reduction of Fe^3+^ to Fe^2+^ ions (Benzie and Strain, 1996). In brief, FRAP reagent (150 μl) was mixed with tissue homogenate (5 μl) and deionized water (15 μl). After incubation for 4 min at 37 °C, absorbance of sample mixture was taken at 593 nm. Aqueous solutions of known Fe^2+^ concentration (0.1-1 mM) was used to generate a calibration curve. Reducing ability of the samples were calculated with reference to the reaction signal from known Fe^2+^ concentration. The values were expressed as micromole of Fe^2+^ equivalents per gram of tissue.

### 2.8 Comet assay

Single strand breaks in the DNA of cortical and hippocampal cells were evaluated by comet assay following earlier described method (Lai and Singh, 1995). In brief, microscopic slides (in duplicate for each sample) were coated with normal melting agarose (NMA, 0.75 %) in deionized water and thoroughly dried at room temperature. Single cell suspension (10 μl) of tissue was mixed with low melting agarose (LMA, 0.5 %, 100 μl) in PBS at 37 °C, spread on pre-coated slides and allowed to solidify for 10 min at 4 °C. Slides were placed in freshly prepared cold lysis buffer (2.5 mM NaCl, 100 mM EDTA, 10 mM Trizma base, 1 % tritonX, 10 % DMSO, pH 10) for 4 hours at 4 °C, then immersed into cold electrophoresis buffer (300 mM NaOH, 1 mM EDTA, pH >13) for 20 min followed by electrophoresis for 25 min at 24 V. After electrophoresis, slides were drained and slowly dipped in neutralization buffer (0.4 M tris, pH 7.5) three times for 5 min each. Next, slides were fixed in chilled ethanol (100 %) for 20 min, and air dried. All the steps were performed under dim light at 4 °C. Slides were randomized and coded before scoring by an independent author. On the day of analysis, slides were rehydrated with ice cold deionized water, stained with ethidium bromide (20 μg/ml, 20 μl) for 2 min and coversliped. Slides were visualized at 40 x magnification under a fluorescent microscope (IX71, Olympus, Japan). Images (n=50) of randomly selected nuclei (25 from each slide) from each animal were analyzed by CometScore software (version 2.0). DNA damage was quantified in terms of percentage of total DNA that is fragmented and migrated away from nuclear DNA (head DNA) forming a comet pattern called tail DNA (%). Whole cell heads without any tail are considered undamaged.

### 2.9 5’-bromo-2’-deox-yuridine (BrdU) administration and cell proliferation assay

To evaluate the effect of RF-EMR exposure on proliferation of NPCs in DG, experimental rats from each group (n=6) were introduced single intraperitoneal injection of 5’-bromo-2’-deox-yuridine (BrdU) (Sigma, USA) at a dose of 300 mg/kg just after the end of exposure period. One set of the animals (n=3) were sacrificed 2 h or 24 h post injection and the effect of RF-EMR precisely on NPCs proliferation were monitored.

Post BrdU administration (2/24 hours) rats were deeply anesthetized with intra peritoneal introduction of sodium pentobarbital (50 mg/kg) and transcardially perfused with cold PBS (0.1 M, pH 7.5) for 5 min followed by cold paraformaldehyde (2 %). Brain was harvested and fixed in paraformaldehyde (2 %) for 48 h at 4 °C followed by immersion in sucrose solutions (10 %, 20 % and 30 %) sequentially to cryoprotect the tissue. Next, brain tissue was embedded into Jung tissue freezing medium (Leica, Germany) and coronal sections were cut (15 μm thickness) through the entire hippocampus (Bregma −1.6 to −6.4) using a Cryostat (Leica CM 1860, Germany). Sections were mounted on gelatin coated slides and were stored at −20 °C. Every 36^th^ section (9-10 section/animal) throughout the hippocampus was processed for BrdU immunofluorescence staining.

For BrdU immunofluorescence staining slides were air dried for 1 h at 37 °C and washed with PBS three times for 5 min each. DNA denaturation was done by placing the slides in 2 M HCl for 15 min at 37 °C. Acid was completely neutralized by three washes of PBS (pH 7.5) for 5 min each. Sections were blocked with normal goat serum (2 %, Merk Millipore, USA) for 1 h. Next, sections were incubated in a humid chamber with mouse monoclonal anti-BrdU (B35128, Invitrogen, USA) primary antibody (1:200 dilution) for 24 h at 4 °C followed by three washes of PBST (0.1 %) for 5 min each. Then, sections were incubated with Alexa Fluor 488-conjugated goat anti-mouse (A11001, Invitrogen, USA) secondary antibody (dilution of 1:500) for 2 h in dark at room temperature. After three washes with PBST (0.1 %) for 5 min each, slides were mounted with fluoroshield mounting medium (Abcam, United Kingdom) supplemented with 4’,6-diamidino-2-phenylindole (DAPI) as counter stain and cover sliped.

Slides were coded and randomized by a co-author other than the experimenter before scoring. All the BrdU-positive cells in the DG (granule cell layer and sub granular zone) were counted under a confocal microscope (FluoView™ FV1000, Olympus, Japan). Counting of BrdU-positive cells was performed at 10 x magnification, while to distinguish single cells within a cluster 100 x magnification was used. The total number of BrdU-positive cells were counted and multiplied by 36 to obtain the total number of cells per DG.

### 2.10 Cresyl violet (CV) staining

Animals (n=3) from each group were sacrificed immediately after the end of exposure. Transcardial perfusion, fixation of brain tissue and sectioning was performed as explained in earlier section. Total five equi-spaced sections per animal spanning somatosensory cortex and hippocampus each were processed for cresyl violet (CV) staining. Briefly, slides with sections were air dried and placed in alcohol chloroform solution (1:1, v/v) for overnight to clear fat followed by rehydration using alcohol (100 %, 95 %) and deionized water sequentially. Sections were stained with CV (0.1 %) staining solution for 5 min at 37 °C and differentiated using ethanol (95 %) for 5 min. Differentiated sections were dehydrated with ethanol (100 %) for 2 min and cleared in xylene for 2 min before cover slipping with DPX mounting medium. Stained sections were visualized using a light microscope (BM-Smart, Lmi Microscopes, England).

Layer IV of somatosensory cortex and different subregions of the hippocampus (DG, CA3 and CA1) were identified and delineated according to the Paxinos and Watson (2006) rat brain atlas. One image from each of the subregions of interest and at equivalent position across sections was captured at 40 x magnification. The number of normal neurons and degenerated neurons in each image (1 image/region/section) were counted using Image J software (http://imagej.nih.gov/ij) by experimenter blind to group codes. Neurons with large and abundant Nissl bodies were considered normal. Degenerated neurons have reduced or no Nissl body and showed heterogenous morphology including nuclear pyknosis, karyorrhectic nuclei, vacuolation etc. Data was presented as percentage of total number of neurons with normal morphology. Values from 5 sections were averaged to use as statistical unit for comparison between the groups.

### 2.11 Statistical analysis

A two-tailed unpaired Student’s t-test was used to compare between the groups and p-value ≤ 0.05 was considered as significant. All results were analyzed using GraphPad Prism software (version 7.01, USA) and expressed as mean ± standard error of mean (SEM).

## 3. Results

### 3.1 Exposure to RF-EMR cause oxidative stress

EPR spectra of cortex and hippocampus tissue homogenate showed dominance of a broad central signal, centered on g=2.0035 (**Fig. 2A**). This value of g-factor denotes presence of carbon centered radicals with adjacent oxygen in the sample (Barclay and Vinqvist, 1994). A comparison of peak to-peak amplitude of radical signal, demonstrated significant increase in g = 2.0035 radical concentration in the cortex (p=0.04) and hippocampus (p=0.047) of RF-EMR exposed rats with respect to sham. Concentration of radical species was increased by ~1.8-fold in cortex region and ~1.7-fold in hippocampus region (**Fig. 2B**).

**Figure 2.**
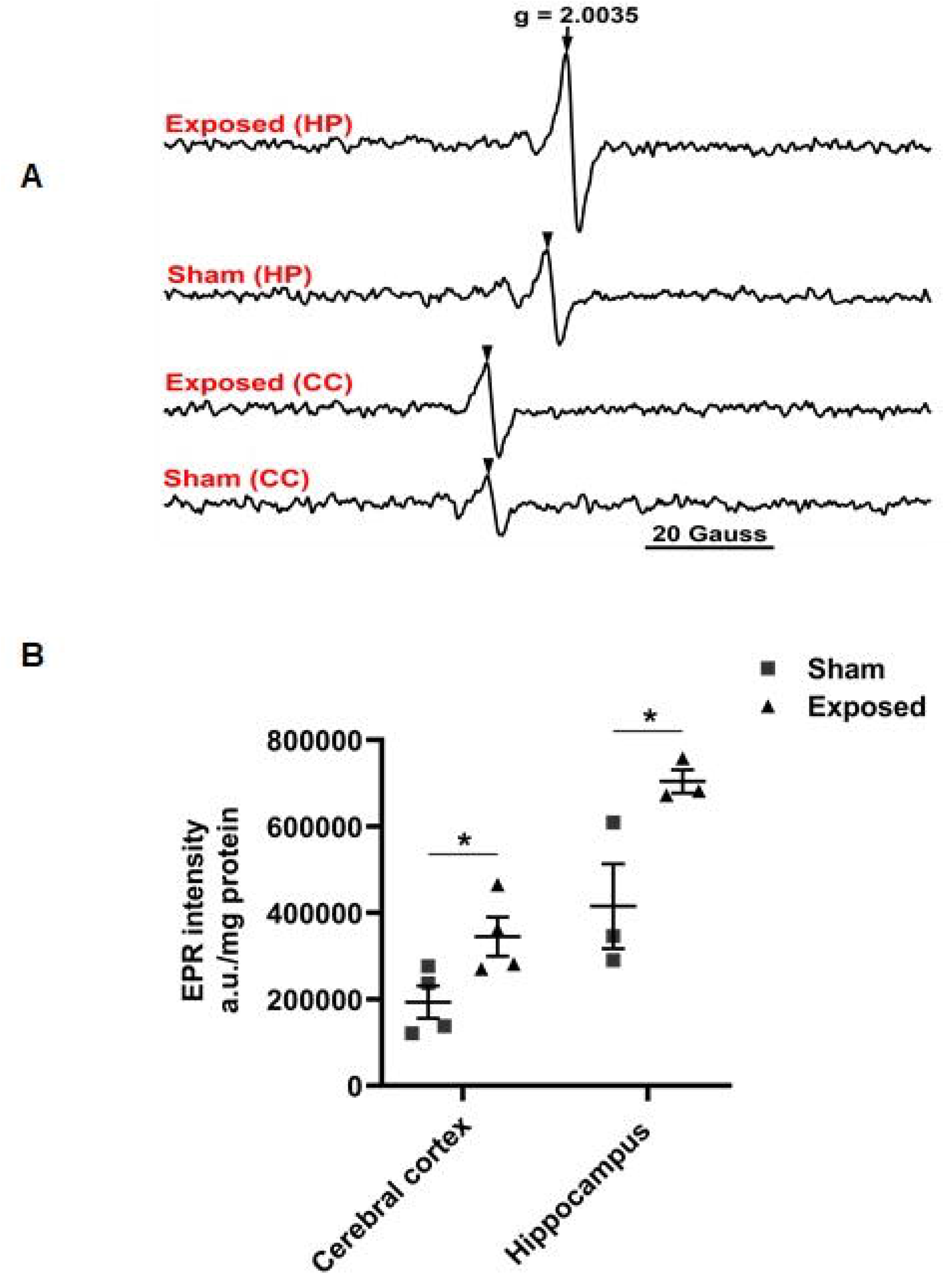
Exposure to radiofrequency electromagnetic radiation (RF-EMR) for 8 hours increased the level of carbon cantered free radicals in the developing rat brain. **A.** Representative images of electron paramagnetic resonance (EPR) spectra showing a clear peak of g = 2.0035 radical species in cerebral cortex (CC) and hippocampus (HP) region of sham and RF-EMR-exposed rats. **B.** Scatter dot plot is showing statistical comparison of EPR signal intensity between groups (n=4). EPR signal intensity was obtained by measuring peak to peak height of central signal which is an estimate of radical concentration and expressed in arbitrary units (a.u.)/mg protein. Data presented as mean ± standard error of mean (SEM). Two-tailed unpaired t-test. *P≤0.05, significantly different from sham group.

Lipid peroxides are the measure of oxidative damage to lipids. A significantly higher level of lipid peroxides was observed in the cortex (p=0.003) and hippocampus (p=0.01) of rat brain exposed to RF-EMR as compared to sham. Lipid peroxides level were increased by 5-fold in the cortex and 3.7-fold in the hippocampus region (**Fig. 3A**). The level of protein carbonyls represents quantity of oxidized protein. No significant alteration in protein carbonyl concentration was observed in either the cortex (p=0.7) or hippocampus (p=0.8) region of rat brain exposed to RF-EMR as compared to shams (**Fig. 3B**). Similarly, the change in antioxidant capacity of cortex and hippocampus region upon RF-EMR exposure was nonsignificant (p=0.3, 0.2) (**Fig. 3C**).

**Figure 3.**
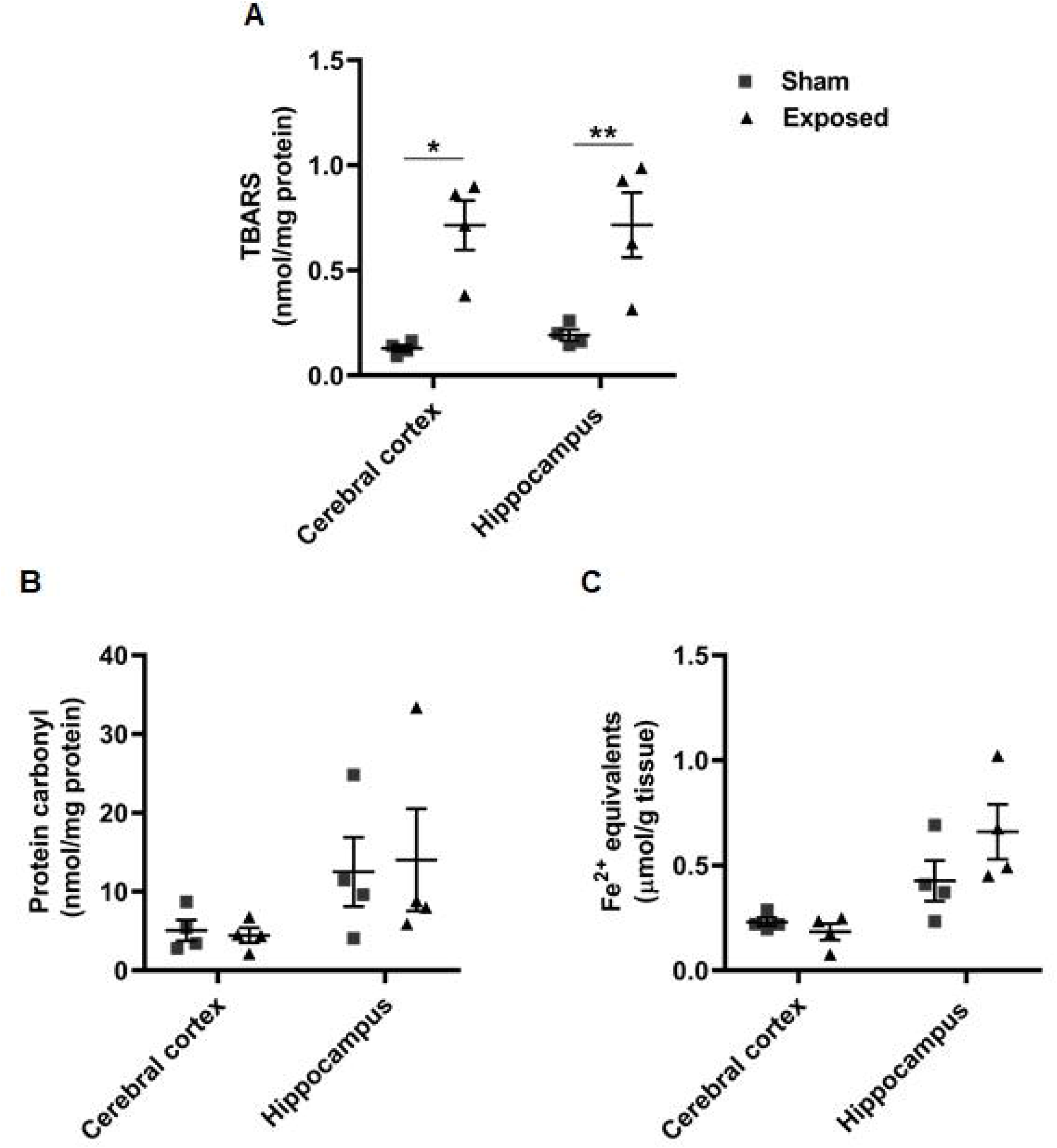
Exposure to radiofrequency electromagnetic radiation (RF-EMR) for 8 hours lead to increased lipid peroxidation in both cerebral cortex (CC) and hippocampus (HP) region of rat brain however, level of protein carbonyl and total antioxidant capacity (TOC) was not changed significantly in either of brain region compared to sham (n = 4). Scatter dot plots are showing statistical comparison between sham and exposed groups for **A.** Lipid peroxidation in nmol TBARS/mg protein **B.** Protein carbonylation in nmol protein carbonyls/mg protein and **C.** Total antioxidant activity in μmol FeSO4 equivalents/g tissue in cortex and hippocampus region of rat brain. Data presented as mean ± standard error of mean (SEM). Two-tailed unpaired t-test. *P≤0.05, **P≤0.01, significantly different from sham group.

### 3.2 Exposure to RF-EMR damages DNA in the cortex and hippocampus

Cells from both cortex as well as hippocampus of rats exposed to RF-EMR showed a comet pattern with clear tail representing migration of fragmented nuclear DNA (**Fig. 4A**). Tail DNA (%) was increased by 2.5-fold in cortex (p=0.001) and 2-fold in hippocampus (p=0.048) of rats exposed to RF-EMR compared to sham animals (**Fig. 4B**). Cortex region seems to be relatively more susceptible to RF-EMR induced DNA damage than hippocampus.

**Figure 4.**
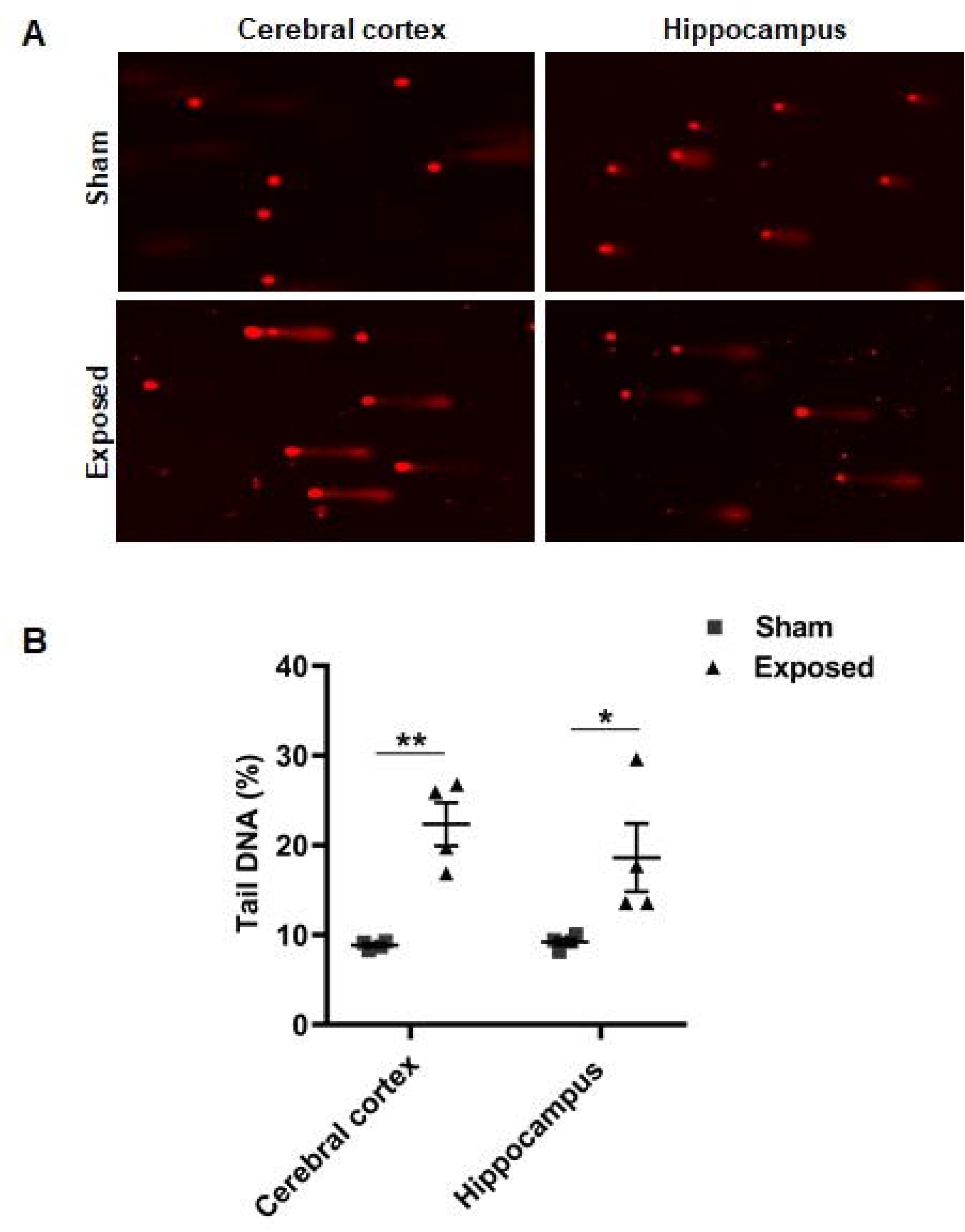
Exposure to radiofrequency electromagnetic radiation (RF-EMR) for 8 hours lead to DNA damage in cerebral cortex (CC) and hippocampus (HP) region of rat brain. **A**. Representative photomicrographs of comets at 10 x magnification from CC and HP region of sham and exposed animals. **B**. Scatter dot plot showing statistical comparison of percentage of total DNA (head DNA), fragmented and migrated to form a comet pattern (tail DNA) between sham and exposed groups (n=4). Whole cell heads without any comet pattern were considered undamaged. For each region tail DNA (%) was estimated in randomly selected 50 individual cells per animal using CometScore software (Version 2.01) and averaged to be used as statistical unit to compare between groups. Data presented as mean ± standard error of mean (SEM). Two-tailed unpaired t-test. *P≤0.05, significantly different from sham group.

### 3.3 Exposure to RF-EMR reduces neurogenesis

We further studied the effect of RF-EMR exposure on hippocampal neurogenesis using BrdU as an endogenous marker. The BrdU-positive cells in the DG were mostly localized in sub granular zone. They were clustered in all animals sacrificed at either 2 or 24 h post BrdU injection, showing diffused patterns of BrdU staining and irregularly shaped nuclei (**Fig. 5A**). Quantification of BrdU-positive cells at both 24 h and 2 h time point demonstrated significantly reduced number of BrdU-positive cells in DG of rats exposed to RF-EMR compared to shams (p=0.03, p=0.049). Number of BrdU-positive cells decreased by 1.6-fold when counted 24 h post injection and 1.4-fold when counted 2 h post injection, suggesting an overall reduced neurogenesis (**Fig. 5B**).

**Figure 5.**
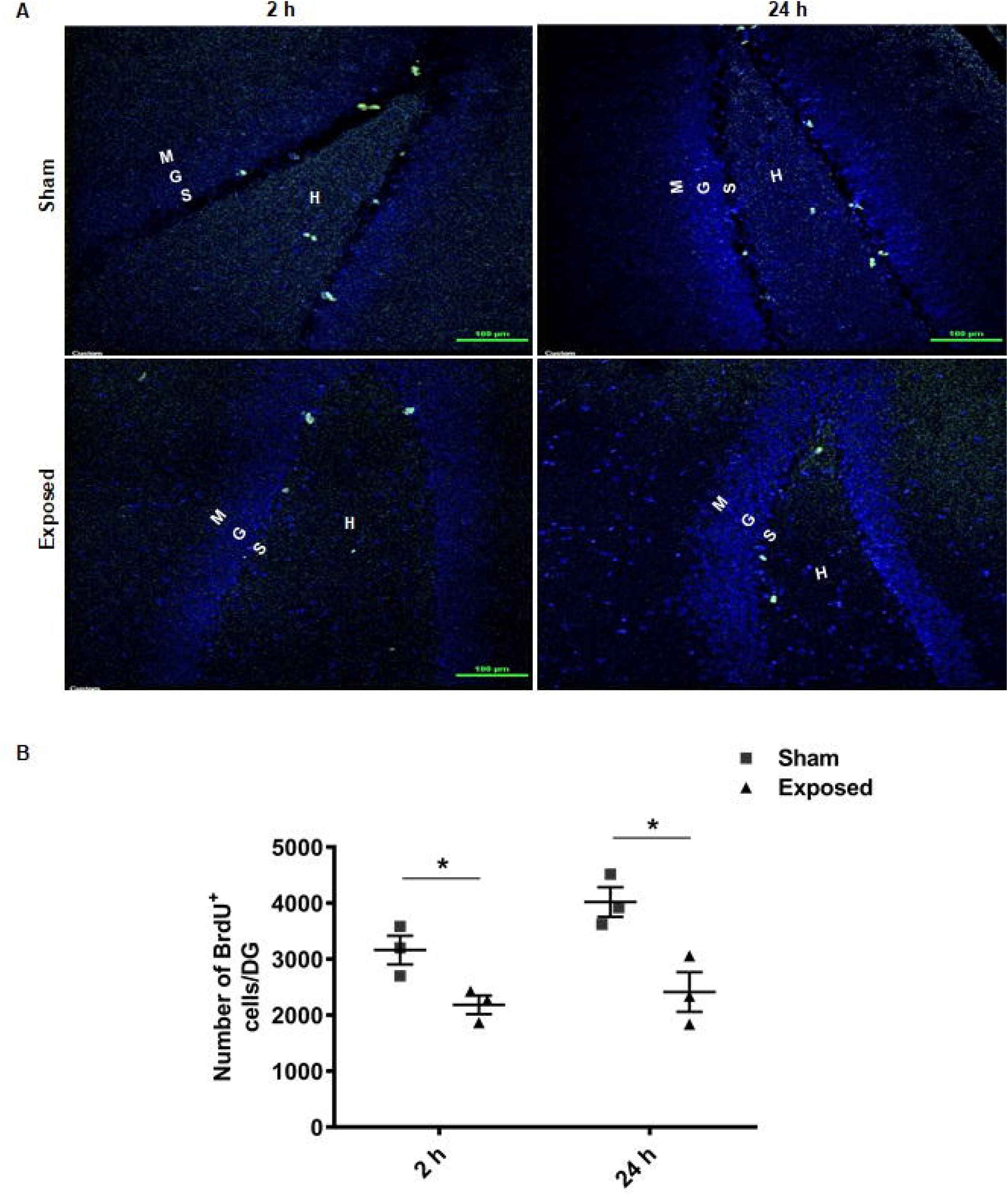
Exposure to radiofrequency electromagnetic radiation (RF-EMR) for 8 hours suppressed hippocampal neurogenesis in Wistar rats. **A**. Representative confocal images showing BrdU-positive cells (green) and counter stained nuclei (blue) in dentate gyrus of sham and exposed rats (Scale bar=100 μm). **B**. Scatter dot plot is showing statistical comparison of BrdU-positive (BrdU^+^) cells between sham and exposed group (n=3) at 2 h or 24 h post injection time points. BrdU-positive cells in the dentate gyrus (granule cell layer and sub granular zone) and hillus were counted in every 36^th^ section and multiplied by 36 to obtain the total number of BrdU-positive cells per dentate gyrus per animal which is used as statistical unit. Data presented as mean ± standard error of mean (SEM). Two-tailed unpaired t-test. *P≤0.05, significantly different from sham group. BrdU: 5’-bromo-2’-deox-yuridine, G: granule cell layer, H: hilus, M: molecular cell layer, S: sub granular zone.

A comparison between number of BrdU-positive cells at 2 h and 24 h time point in the same group, provides information about the number of cells normally completing the cell cycle. It was found that DG region of sham animals have 3163 ±443 BrdU-positive cells at 2 h and 4021 ± 456 cells at 24 h time point amounting to 1.3-fold increase in number, whereas, DG region of rats exposed to RF-EMR have 2187 ±289 BrdU-positive cells at 2 h time point and 2413 ± 413 cells at 24 h time amounting to 1.1-fold increase in number. These results suggest that RF-EMR exposure might also be interfering with the cell cycle progression in NPCs (**Fig. 5B**).

### 3.4 Exposure to RF-EMR causes neuronal degeneration in DG

We further examined the effect of RF-EMR exposure on morphology of neurons in cerebral cortex, and subregions of hippocampus (DG, CA3 and CA1) (**Fig. 6A**). Degenerating neurons were detected and quantified using Nissl staining. We observed neurons with abnormal morphology (pyknotic or karyorrhectic nuclei, vacuolization) in all of these regions of rat brain exposed to RF-EMR, however the changes were more prominent in the DG region of hippocampus (**Fig. 6B**).

**Figure 6.**
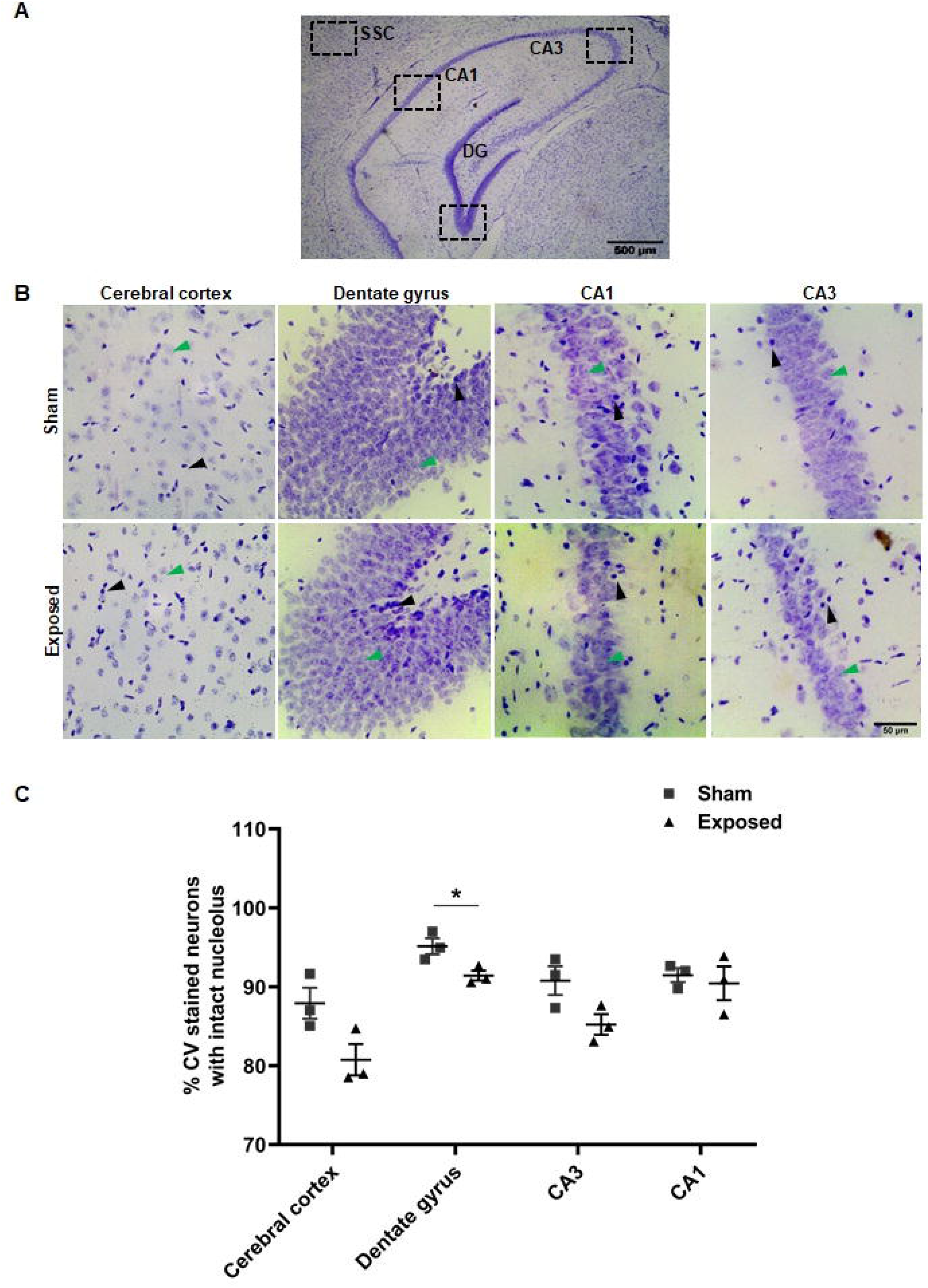
Exposure to radiofrequency electromagnetic radiation (RF-EMR) for 8 hours lead to morphological changes in dentate gyrus (DG). **A.** Photomicrograph of a cresyl violet (CV) stained coronal section (Scale bar=500 μm) of rat brain showing somatosensory cortex and different subregions of hippocampus (CA1, CA3, and DG). Respective regions used for analysis were further delineated in boxes. **B.** Representative photomicrographs of CV stained sections (Scale bar = 50 μm) of sham and exposed rats. Black and green arrow shows pyknotic and healthy neurons respectively. **C.** Scatter dot plot is showing statistical comparison of percentage of intact neurons in cerebral cortex, and hippocampal subregions (dentate gyrus, CA1 and CA3) between sham and exposed groups (n =3). One field of view from equivalent position across animals for each region and each section was analyzed using Image J software to count total number of healthy and pyknotic neurons. The mean cell count (presented as percentage of healthy neurons) from five sections per animal was used as statistical unit. Data presented as mean ± standard error of mean (SEM). Two-tailed unpaired t-test. *P≤0.05, significantly different from sham group. CA: Cornu ammonis, SSC: somatosensory cortex.

Quantitative analysis showed a significant decrease in the percentage of neurons with normal morphology (large and abundant Nissl’s bodies) in the DG of rats exposed to RF-EMR as compare to shams (p=0.03). The morphological changes were insignificant in CA3 (p=0.07) and CA1 (p=0.7) subregions of hippocampus and cerebral cortex (p=0.06) (**Fig. 6C**). Based on these results, neurons in the DG region appeared to be more vulnerable to RF-EMR exposure compared to neurons in the other regions.

## 4. Discussion

Extensive use of mobile phones by children and young adolescents in recent decade has been reported to be associated with behavioral and cognitive disorders (Foerster et al., 2018). The underlying mechanisms behind these changes are not clearly understood. The present study showed that exposure of 5-week old male Wistar rat to 2115 MHz RF-EMR for 8 h lead to oxidative stress and DNA damage in cortex and hippocampus region of brain along with reduced neurogenesis and neuronal degeneration in DG region of hippocampus (**Fig. 7**). These results augment our understanding to evaluate the effect of RF-EMR on young adolescent brain which is undergoing critical developmental changes and maturational process.

**Figure 7.**
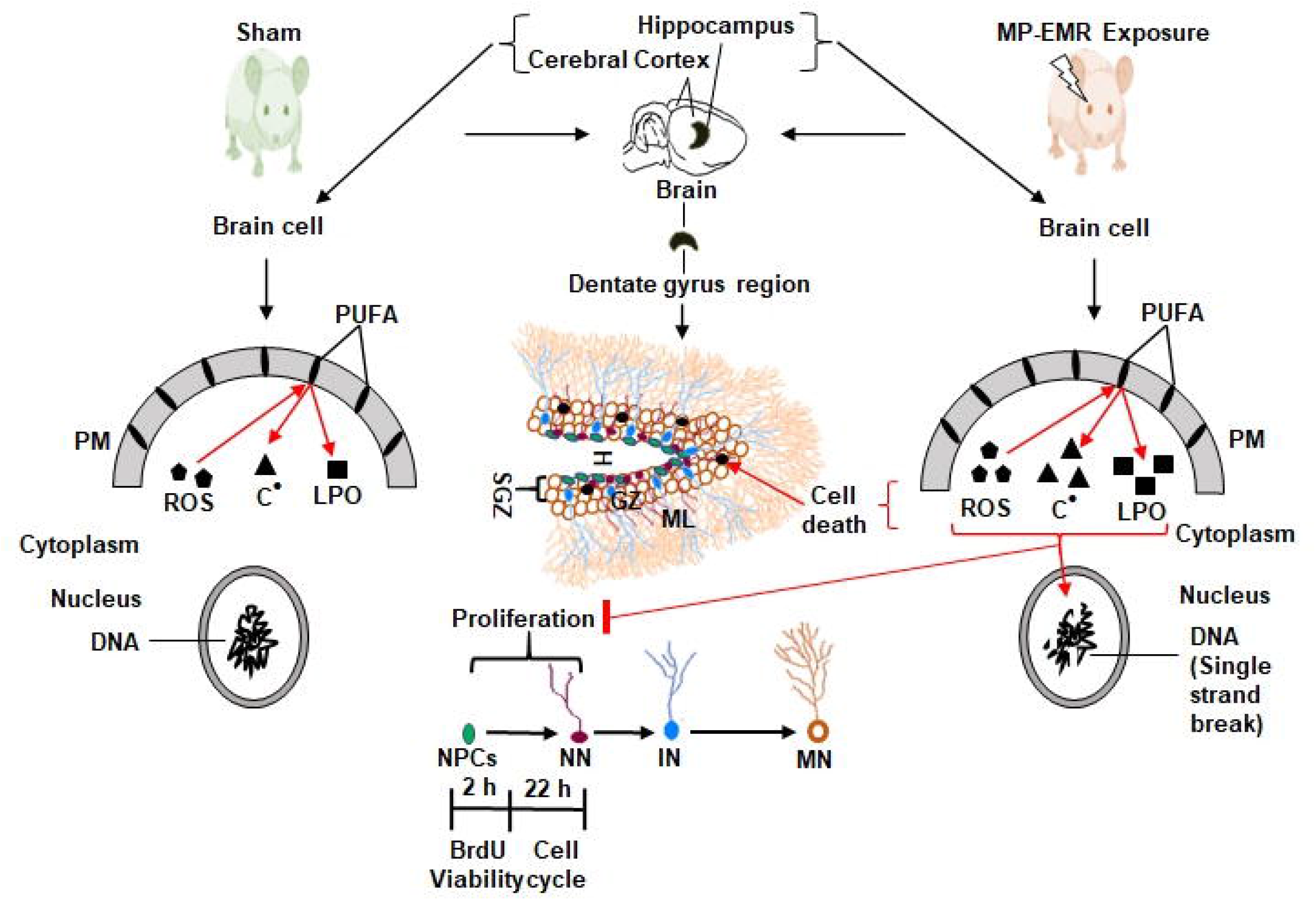
Exposure to radiofrequency electromagnetic radiation (RF-EMR) seems to induce reactive oxygen species (ROS) production which targets polyunsaturated fatty acids (PUFA) in cerebral cortex and hippocampus cell membrane and generates carbon centered radicals (C) and lipid peroxides (LPO) leading to DNA damage, neuronal degeneration and impaired proliferation of hippocampal neural progenitor cells (NPCs). Important outcomes and their possible role and interrelationship at cellular level are presented. BrdU: 5’-bromo-2’-deox-yuridine, GZ: granular zone, H: hillus, IN: immature neurons, ML: molecular layer, MN: mature neurons, NN: new born neurons, PM: plasma membrane, RF: radiofrequency, SGZ: sub-granular zone.

We found a significant increase in concentration of free radicals, especially carbon center radicals and lipid peroxides in cortex and hippocampus regions of rat brain upon RF-EMR exposure. However, the protein carbonyls and total antioxidant capacity has not shown significant change in either of brain regions of exposed rats. Previously, various animal models and cell line studies have reported that RF-EMR exposure produces ROS (Yakymenko et al., 2016). Generation of RF-EMR induced-ROS in plasma membrane seems to be mediated by NADH oxidase (Friedman et al., 2007), a key component of mitochondrial electron transport chain. Once produced, these ROS in turn attacks polyunsaturated fatty acids (PUFA) of cell membrane and form transient carbon centered lipid radicals. These lipid radicals react with the molecular oxygen forming lipid peroxyl radical which ultimately decomposes in to a variety of toxic end products like aldehydes and ketones (Aruoma, 1998; Marnett, 1999).

Brain tissues have high membrane-to-cytoplasm ratio (Evans, 1993) and in the membrane PUFA are predominant lipids (Watson, 1993). This might be the probable reason that in this study the effect of RF-EMR exposure is more pronounced on carbon centered radicals and lipids than on other oxidative stress markers. Elevated level of radical species can also oxidize proteins and reduces the antioxidant reservoir of cells (Bayir et al., 2002). Earlier reports have demonstrated that RF-EMR exposure causes lipid peroxidation, protein carbonylation and diminish cellular antioxidant defense (Alkis et al., 2019; Megha et al., 2015). In these studies, repeated RF-EMR exposure was provided for a long period of time, however, we assessed immediate effects of a single acute exposure of RF-EMR, which might be the reason that we did not observed significant changes in the level of protein carbonyls and total antioxidant capacity.

Once produced inside the cells, different radical species including lipid derived radicals may cause oxidative damage to cellular macromolecules including nucleic acid (Marnett, 1999; Belpomme et al., 2018). In the present study, we observed single strand DNA break along with increased carbon centered lipid radicals and lipid peroxides in the cortex and hippocampus region of rat brain exposed to RF-EMR. Earlier reports showed mixed results on DNA damage in brain after exposure to varying doses and durations of RF-EMR (Belyaev, 2015; Prihoda, 2019). A number of studies demonstrated DNA damage after 2 h of 2450 MHz radiation exposure as well as 4 h post exposure at a SAR value of 0.6 or 1.2 W/Kg (Li and Singh 1995, 2005). Lagroye et al. (2004) observed no DNA damage on same exposure parameters. Similarly, chronic microwave exposure studies also reported increased DNA damage in a frequency dependent manner with maximum damage at 2450 and 2100 MHz frequencies compared to 900 or 1800 MHz (Alkis et al., 2019; Deshmukh et al., 2015; Megha et al., 2015), while few studies didn’t observe significant DNA damage irrespective of frequencies (Su et al., 2018; Verschaeve et al., 2006). Most of these studies were performed on adult brain and indicates that frequency, dose and duration of exposure might be critical for the extent of DNA damage. In this study, rats age matched to young adolescents were exposed to comparatively higher microwave frequency (2115 MHz) and SAR for 8 h, that may be the reason we observed notable DNA damage in brain cells of exposed animals. RF-EMR induced DNA damage might be the consequences of high amount of carbon centered free radicals produced under oxidative stress in brain. The DNA damage was more pronounced in cortex region than hippocampus owing to its proximity to source of radiation (Cardis et al, 2008), and may have absorbed higher RF radiation.

Radical species-mediated oxidative damage has reported to impair neurogenesis and cell survival which is associated with increased risk of neurological disorders (Bazinet and Laye, 2014). BrdU is a thymidine analogue which labels all actively dividing cells in S-phase. In present study, we observed reduced number of BrdU-positive cells in DG of rats exposed to RF-EMR compared to shams at both 2 h or 24 h post BrdU injection time points. Two different time points (2 h and 24 h post BrdU injection/irradiation) were chosen, as BrdU has bioavailability for 2 h, while 24 h is the approximate cell cycle time of NPCs (Cameron and Mckay, 2001). Reduced number of BrdU-positive cells at these time points indicates that RF-EMR exposure may have either inhibited cell division or have lengthened cell cycle duration or interfered with cell cycle progression in NPCs of DG. Moreover, in exposure group, only 55 % of NPCs which are initially labeled (2 h time point) have completed the cell cycle (24 h time point) amounting to 1.1-fold increase in number of BrdU-positive cells. However, under normal condition, for rats of this age group, about 75 % of S-phase NPCs complete the cell cycle during this time period, amounting to 1.5-fold increase in number of BrdU-positive cells (Cameron and Mckay, 2001). Hence, it seems that RF-EMR exposure may have interrupted progression of cell cycle in NPCs of DG region. However, to ascertain the exact stage at which RF-EMR has interrupted the proliferation of NPCs, further studies are needed. Earlier studies investigating the effect of RF-EMR exposure on adult neurogenesis, precisely in DG region of hippocampus are very few with mixed findings. Exposure of 1.8 GHz RF-EMR at SAR of 1.16 W/kg for 8 h for 3 consecutive days stimulated DNA synthesis but interrupted cell division and stem cells pool in DG of new born rats while showed no effect on juvenile brain (Xu et al., 2017). Similarly, mice exposed to 2.45 GHz microwave frequency for 2 h and sacrificed 2 h post irradiation has shown reduced neuronal proliferation in subventricular zone of new born but no effect in young ones (Orendacova et al., 2011). However, Kim et al. (2015) found no effect of 848.5 MHz CDMA signal on the number of BrdU-positive cells in DG of young adult rats after 2 weeks of exposure. Similarly, number of PCNA positive cells (an early marker of neuronal proliferation) in subventricular zone and DG of adult mice exposed to RF-EMR at SAR of 7.8 W/kg for 6 or 12 months (1 h daily, 5 days/week) remain unaffected (Kim et al., 2008). These studies provide insight on the effect of RF-EMR on neurogenesis and shows its dependence on developmental stages of brain and duration of exposure. Furthermore, these studies suggest that younger brain and short-term RF-EMR exposure seem to have greater effect on neurogenesis which corroborates our findings.

Histopathological observations of neurons present an idea about neuronal health and degeneration. We found markedly increased neuronal degeneration in DG of hippocampus whereas other sub regions like CA3, CA1 and cerebral cortex have not shown remarkable changes. The morphological alterations observed in neuronal cells have clearly shown the characteristic of their degeneration, and indicated that RF-EMR has affected the survival of proliferating NPCs more, than the well differentiated neurons. This might be possible, as developing brain has large number of NPCs in DG, undergoing proliferation and differentiation (Qiu et al., 2007) and are highly susceptible to oxidative damage and apoptosis under any stress situation including RF-EMR exposure (Leone et al., 2014; Naylor et al., 2008). Most of the damaged neurons have large vacuoles and are interspersed between normal neurons indicating damage happened in the living cells. These findings are in agreement with previous reports demonstrating neuronal degeneration in different brain regions after microwave exposure of varying frequencies, doses and durations (Bas et al., 2009; Odaci et al., 2008; Saikhedkar et al., 2014; Tan et al., 2017). Salford et al. (2003) demonstrated prominent neuronal damage in cortex, hippocampus and basal ganglia by nonthermal microwave exposure of very low GSM signals (at a SAR of 0.002, 0.02 and 0.2 W/Kg) only for 2 h and sacrificed 50 days post irradiation. Present study provide evidence regarding specific vulnerability of neurons in a specific region of developing brain towards apoptosis upon acute exposure to RF-EMR.

## 5. Conclusion

In conclusion, this study exhibited that RF-EMR exposure for long time induces oxidative stress in different regions of young adolescent rat brain with marked increase in carbon centered radicals and lipid peroxidation. The cortex and hippocampus of rats exposed to RF-EMR, showed DNA damage in the form of single strand breaks, impaired hippocampal neurogenesis and increased neuronal degeneration in DG region. Despite the fact that studies in animal model can’t be directly extrapolated to humans, combined results of our study showed adverse effect of high frequency RF-EMR in rat brain. These findings after careful validation may be useful to develop appropriate interventions for reducing the impact of RF-EMR induced adverse effects in humans.

## Appendix A: Supplementary material

**Supplementary Fig. 1** Image of experimental set up used to expose Wistar rats to RF-EMR for 8h. **Supplementary Table 1** Parameters of RF-EMR exposure.

## Author contributions

KVS, RKN and PR made substantial contribution to conception and design, data acquisition, analysis, and interpretation. KVS conducted the experiments. CP contributed in performing immunohistochemical experiments. JPN contributed in conducting exposure experiments and dosimetry. KVS wrote the original draft. KVS, CP, PR and RKN edited the manuscript. RKN and PR acquired the funds and supervised the overall research work.

## Ethical Approval

The experimental procedures were approved by the institutional animal ethical committee (IAEC) of Jawaharlal Nehru University, New Delhi, India (Application code # 23/2016). Experiments were performed in accordance with the committee for the purpose of control and supervision of experiments on animals (CPCSEA) guidelines.

## Acknowledgement

This work is partly supported by UPE II project (ID-58) and DST PURSE fund of Jawaharlal Nehru University (JNU) and institute core funds from the International Center for Genetic Engineering and Biotechnology (ICGEB). KVS was supported by a fellowship from Council of Scientific and Industrial Research (Award No. 09/263(1005)/2013-EMR-1) and Indian Council of Medical Research, Government of India (Award No. 849/2019); CP was supported by ICMR-RA fellowship from ICMR, Government of India.

## Abbreviations

BrdU: 5’-bromo-2’-deox-yuridine
CA: Cornu ammonis
CV: Cresyl violet
DG: Dentate gyrus
NPCs: Neural progenitor cells
RF-EMR: Radiofrequency electromagnetic radiation
ROS: Reactive oxygen species
SAR: Specific absorption rate

