## Supplementary Fig. 1 and Supplementary Table 1 for "Acute radiofrequency electromagnetic radiation exposures cause neuronal DNA damage and impair neurogenesis in the young adolescent rat brain"

or,

Ranjan Kumar Nanda (PhD)

Group Leader

Translational Health Group

International Centre for Genetic Engineering and Biotechnology (ICGEB)

New Delhi Component, Aruna Asaf Ali Marg

INDIA-110067

**Contents**

**Supplementary Fig. 1** Image of experimental set up used to expose Wistar rats to radiofrequency electromagnetic radiation (RF-EMR) for 8h.

**Supplementary Table 1** Parameters of radiofrequency electromagnetic radiation (RF-EMR) exposure.

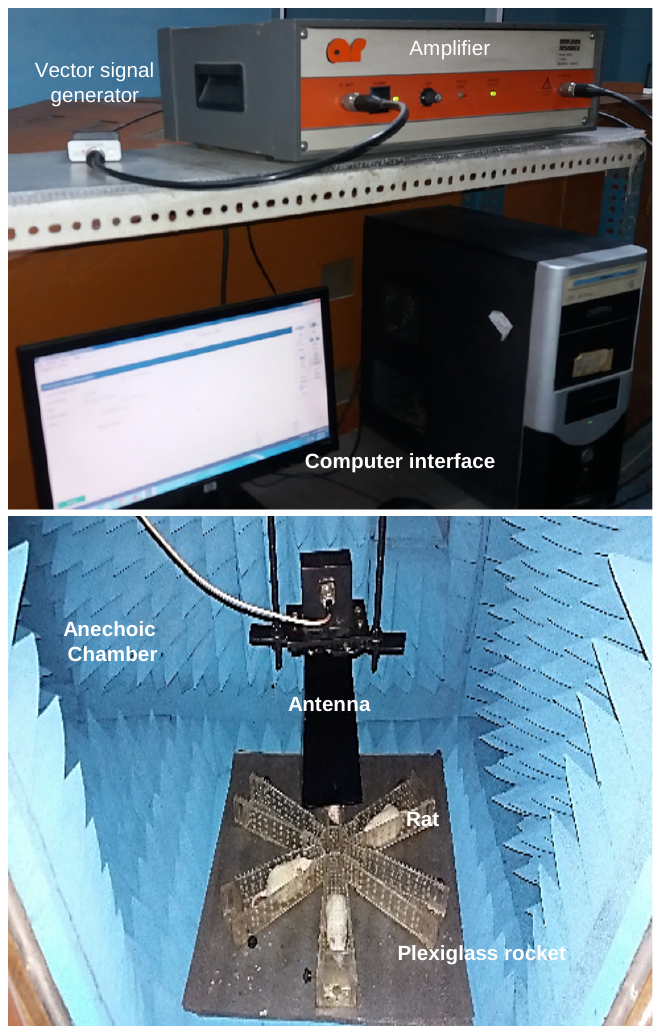

**Supplementary Fig. 1** Young adolescent (5-week age) male Wistar rats were subjected to radiofrequency electromagnetic radiation (RF-EMR) for 8 h in a specially designed exposure setup. RF-EMR radiation of 2115 MHz frequency was generated using vector signal generator attached to an amplifier for power modulation and transmitted through a horn antenna placed inside anechoic chamber. While exposing rats were placed inside plexiglass rockets and kept just below the antenna at the base of anechoic chamber to allow equal exposure of all animals.

**Supplementary Table 1** Parameters of radiofrequency electromagnetic radiation (RF-EMR) exposure measured in terms of frequency, and power density. Whole body averaged specific absorption rate (SAR) was calculated empirically.

| **Exposure Parameters** | |
| --- | --- |
| Frequency | 2115 MHz |
| Power density (off condition) | 0.012 mW/cm^2^ |
| Power density (on condition) | 1.11 mW/cm^2^ |
| Power attenuation (due to 1 mm thick plexi-glass sheet) | 0.11 mW/cm^2^ |
| Power density (net) at the surface of head | 1 mW/cm^2^ |
| Whole body average SAR value | 1.15 W/kg |
